# What is the best fitness measure in wild populations? A case study on the power of short-term fitness proxies to predict reproductive value

**DOI:** 10.1101/2021.11.19.469339

**Authors:** Vita Živa Alif, Jamie Dunning, Heung Ying Janet Chik, Terry Burke, Julia Schroeder

## Abstract

Fitness is at the core of evolutionary theory, but it is difficult to measure accurately. One way to measure long-term fitness is by calculating the individual’s reproductive value, which represents the expected number of allele copies an individual passes on to distant future generations. However, this metric of fitness is scarcely used because the estimation of individual’s reproductive value requires long-term pedigree data, which is rarely available in wild populations where following individuals from birth to death is often impossible. Wild study systems therefore use short-term fitness metrics as proxies, such as the number of offspring produced. This study obtained three frequently used short-term proxies for fitness obtained at different offspring life stages (eggs, hatchlings, fledglings and recruits), and compared their ability to predict reproductive values derived from the genetic pedigree of a wild passerine bird population. We used twenty years of precise field observations and a near-complete genetic pedigree to calculate reproductive success, individual growth rate and de-lifed fitness as lifetime fitness measures, and as annual de-lifed fitness. We compared the power of these metrics to predict reproductive values and lineage survival to the end of the study period. The three short-term fitness proxies predict the reproductive values and lineage survival only when measured at the recruit stage. There were no significant differences between the different fitness proxies at the same offspring stages in predicting the reproductive values and lineage survival. Annual fitness at one year old predicted reproductive values equally well as lifetime de-lifed fitness. However, none of the short-term fitness proxies was strongly associated with the reproductive values. In summary, the commonly short-term fitness proxies capture long-term fitness with intermediate accuracy at best, if measured at recruitment stage. As lifetime fitness measured at recruit stage and annual fitness in the first year of life were the best proxies of long-term fitness, we encourage their future use.

## Introduction

The concept of fitness is central to evolutionary theory (1). Natural selection maximises fitness, which is therefore a driving force of evolution as well as a measure of evolutionary success (2). One definition of fitness is how good an individual is at spreading its genes into future generations, relative to all other individuals in the population (2). A universal definition of fitness in mathematical terms that applies to all population structures and dynamics is however not agreed on (2–5).

Ecological studies measure fitness in diverse ways, often depending on the research question, the population dynamics, and the ecology of the study species (6,7). While some studies measure fitness across lifetimes, other studies measure individual annual fitness to examine variation in selection between years (8). Lifetime fitness is considered more accurate than annual measures, as the latter is influenced by environmental stochasticity (7,9). Alternative fitness measures have been developed that account for environmental stochasticity and population dynamics (5,10–12).

Some fitness metrics include both survival and fecundity components (8,13), while others focus on only one component as a proxy, such as lifespan (4), or on only a single life-history trait, such as the age at first reproduction (13,14). The two most commonly used individual fitness proxies are lifetime reproductive success (LRS) (15) and individual growth rate (IGR) (13). Both count the number of offspring produced in the individual’s lifetime, but IGR gives more weight to offspring produced at a younger age (13), therefore results differ (16). As a consequence, different fitness proxies do not necessarily correlate well (7) and more research is needed to determine which is the most appropriate measures of fitness (6). Choosing the appropriate fitness proxy is therefore an important consideration when designing a study (7).

The age of an offspring at the time it is included in the fitness measurement of the parent must also be considered, particularly given that studies count offspring at varying ages or life-history stages. Offspring survival can be both a part of offspring fitness due to its unique genotype, or a part of parental fitness in cases where parental phenotype affects offspring mortality, for example through parental care (17). Counting offspring at higher stages of development assigns a part of offspring fitness into parental fitness, thus potentially affecting the strength and direction of selection. Furthermore, Brommer et al. (6) show that the age of offspring census directly affects IGR values, while it does not affect LRS, making the two fitness proxies less comparable at different stages. It is therefore important to understand what census time best captures parental fitness.

Although fitness is considered to be a measure of an individual’s gene copy frequency in future generations, most fitness proxies focus on an individual’s direct descendants. Alternatively, the reproductive value from a single individual, defined as the expected number of copies of each of an individual’s alleles in a future generation conditional on a realised pedigree of descendants, can be used to measure long-term fitness (18). The reproductive values can be estimated from a genetic pedigree, following rules of mendelian inheritance to calculate how many allele copies survive on average. The reproductive values stabilise after log_2_*N* generations, where *N* is the population size (4,18,19). While the ultimate genetic contribution of an individual will only emerge over long timescales (>100 generations), the reproductive values are determined in ~10 generations and are a good predictor of the ultimate genetic contribution (18).

The reproductive values closely predict allele survival probability, but not their frequencies (18). Due to recombination and segregation in meiosis, the actual genetic frequencies, conditional on allele survival, instead follow a random distribution (18–21). Consequently, not all genealogical ancestors are also genetic ancestors (22). Despite the difference between actual allele frequencies and the reproductive values, reproductive value is a practical and relevant measure for evolutionary studies as it is maximised by natural selection, thus closely corresponding to fitness (18,23).

This study examined the correlation between several short-term fitness proxies and reproductive values. We used data from an isolated house sparrow (*Passer domesticus*) population on Lundy Island (United Kingdom) with 20 years of life history data, unusually precise measures of survival and reproductive success, and nearly complete genetic pedigree information (24). We examined the two most commonly used individual fitness proxies based on fecundity (25): lifetime reproductive success (LRS) and individual growth rate (IGR) (13). We measured both at four different offspring stages (eggs, hatchlings, fledglings, and recruits) to investigate which are most accurate. We also used a short-term fitness proxy that incorporates survival – de-lifed fitness (8). This is based on individual offspring production and survival adjusted for population growth.

## Methods

### Study system

The house sparrow population on Lundy Island has been continuously monitored since 2000. Lundy is a small island 19 km off the south-west English coast (51°11N, 4°40W). In 2000, 50 individuals were brought to Lundy from the mainland for an experiment (26). Due to the distance of the island from the mainland and the sedentary nature of sparrows, there is minimal dispersal to and from the island (24). The sparrow population size has fluctuated between 166 and 1242 individuals (juveniles included) between 1999 and 2019 (Fig 1A).

**Fig 1:**
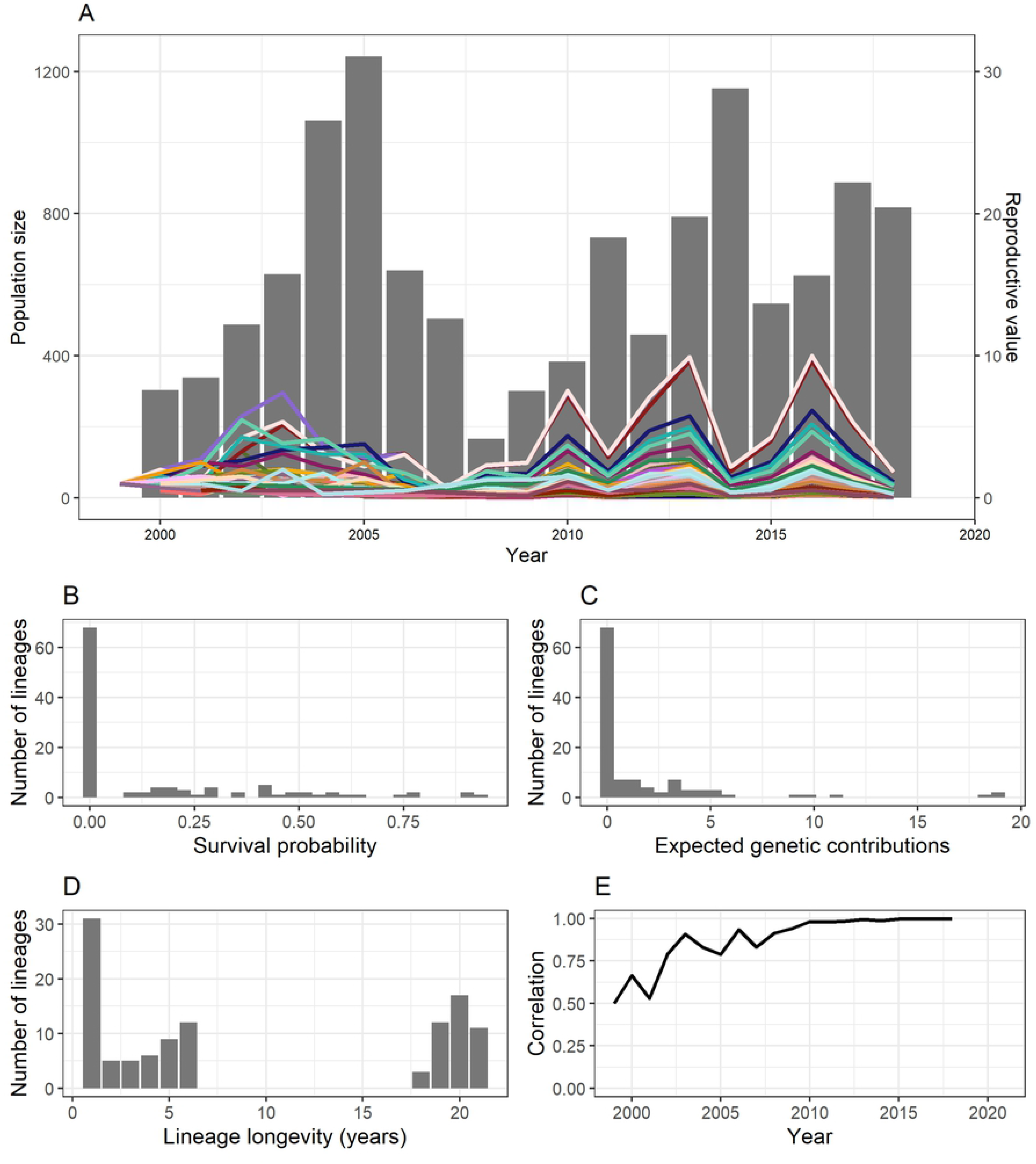
(A) Reproductive values and population size during the study period. Bars represent the population size and lines represent the reproductive values of each of the 43 lineages that survived to 2018. (B) Survival probability of 111 house sparrow lineages on Lundy Island from 1999 to 2018. (C) Reproductive values of the 111 lineages. (D) Number of years to lineage extinction for the 111 lineages. Lineages that survived 18, 19, 20 or 21 years are those that were still extant in 2018, corresponding to cohorts 1999, 2000, 2001 and 2002 respectively. (E) Correlation between the reproductive values in each year and the final year.

During systematic annual monitoring, each sparrow is ringed with three colour rings and one metal ring from the British Trust for Ornithology. Since most sparrows are initially caught as nestlings and ringed as fledglings, we know the identities of the parents attending their nests, and the exact age of all individuals (27). Over 99% of the population has been ringed since 2000 (27). If an individual is not seen for two years or more, it is assumed dead, with this assumption based on previous mark–recapture success data (27,28). Blood samples are collected upon bird capture and genotyped at up to 23 microsatellite loci (24). This allows for the assignment of genetic parentage with 95% confidence (24). From the genetic pedigree and the social brood information, the reproductive success of individuals is calculated. Thanks to these data, the study system provides unusually accurate survival, reproduction, and pedigree data for the complete population (24).

### Pedigree analysis

We calculated fitness proxies and the reproductive values for founders and half-founders that were born between 1999 and 2002, the starting years of the long-term study. Founders and half-founders were defined as individuals for which both parents, or one parent, respectively, were unknown. To calculate reproductive values, we used our genetic pedigree containing all reproducing individuals up to 2018. We removed any individuals from cohorts after 2002 that had at least one unknown parent; thus 8% of all individuals in the pedigree were removed.

Reproductive values were calculated using gene dropping (29). Gene dropping is a computer simulation in which each individual is assigned two alleles (one paternal and one maternal), and their Mendelian transmission down the pedigree is simulated. By repeating this simulation many times and calculating the mean values, robust estimates of reproductive values can be obtained by examining the frequency of an individual’s alleles in subsequent generations. In addition, the allele survival probability can be calculated by examining in how many simulations the allele survives in present-day individuals. We ran the simulation 10,000 times using R package nadiv (30). Using the results, we derived lineage longevity, reproductive values, and allele survival probability. We define lineage longevity as the number of years before a lineage originating from one individual goes extinct, and allele survival probability is the proportion of gene dropping simulations in which a lineage survives. We explored whether lineages from the experimentally introduced sparrows differed from native lineages in their rate of survival to 2018 (last year with complete data) and in their reproductive values. We chose to work with years rather than generations as a measure of time because sparrows have overlapping generations.

### Short-term fitness metrics

We calculated the short-term fitness proxies for the founders and half-founders from cohorts between 1999 and 2002 with complete life-history data. Founders with incomplete life-history data were removed because this could lead to an underestimation of their reproductive success. The individual lifetime production of eggs, broods, hatchlings, fledglings, and recruits was then calculated, as well as IGR at all four offspring stages, and de-lifed fitness. Hatchlings were defined as offspring counted in a nest two days after hatching, and fledglings were birds that survived until ringing, which is typically 12 days after hatching. Recruits were defined as offspring that successfully reproduced and produced at least one egg in any subsequent years.

The IGR is the dominant eigenvalue of an individual population transition matrix, as described in (13). In an individual population transition matrix, the sub-diagonal represents survival, and the first row is filled with the number of offspring produced at each parental age, divided by two to account for parent–offspring relatedness being ½. An example of an individual population transition matrix for an individual that survived three years and had 1, 2 and 1 offspring at ages 1, 2, and 3 respectively, is given below:

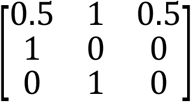

We also calculated annual de-lifed fitness based on the formula (8):

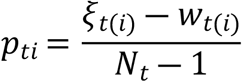

Where:

- *p_ti_* is individual fitness
- *ξ_t(i)_* is the number of individual’s surviving offspring at the end of the time step plus one if the individual survived
- *w_t(i)_* is the population size in year *t*+1 divided by population size in year *t*
- *N_t_* is the population size of adults on 1^st^ April each year

While LRS and IGR are both lifetime fitness measures, de-lifing was designed primarily as a per-generation fitness proxy and is here calculated annually. However, lifetime de-lifed fitness can be obtained by summing the annual fitness values for each individual (8,31). We therefore used de-lifed fitness as both an annual fitness proxy and, after summing, as a lifetime fitness proxy.

We calculated Pearson’s correlation between each fitness metric and the reproductive values for the lineages that survived. We ran a binary logistic regression model in R version 4.0.3 (32), using MCMCglmm (33) with lineage survival to 2018 as the response variable, and each fitness metric as the explanatory variable. The fitness metrics were z-transformed so that the slopes were not affected by the variable variances. We used priors with the residual variance fixed at 0.5 and ran the model for 100,000 iterations with the thinning interval set at 70 and the burn-in at 7,000. We examined which fitness metric had the strongest association with reproductive values based on the slope of the regression.

### Ethics statement

As this was a theoretical study using previously selected data, no ethics approval was required.

## Results

### Reproductive values

There were in total 111 lineages arising between 1999 and 2002 used for the analyses. Of these 111 lineages, 18 lineages were founded by sparrows experimentally introduced in 2000 (26) and 93 lineages stem from native sparrows already present on the island in 2000. Forty-three lineages survived to 2018, of which 11 were introduced and 32 were native. Hence, at most, 39% of the founders passed genetic material to 2018, and there was no statistically significant association between a lineage’s origin and survival (*p* = 0.06, Fisher’s exact test). The mean lineage survival probability was 0.16 (95% CI 0.13–0.18), and the survival probability for lineages appearing in 2018 was 0.40 (95% CI 0.37–0.43, Fig 1A). There was variation in the absolute reproductive values (mean = 1.64, 95% CI 1.32–1.95 range: 0.41–18.89, Fig 1B). The introduced lineages had on average higher reproductive values than native lineages (*t* = 2.70, df = 17.90, *p* = 0.015). Contributions varied over time, but after 2007 fluctuations were more synchronous among lineages, and the ranking of lineages based on their reproductive values remained similar (Fig 1E). Population fluctuations closely follow fluctuations in the reproductive values in the previous year. The change in lineage behaviour after 2007 is visible in lineage longevity, as all lineages that survived the from 2000 to 2006 also survived until 2018 (Fig 1C). After 2007, the correlation between annual reproductive value and reproductive values in 2018 also increased, and stabilised around 2011 subsequent to which the correlation was above 0.95 (Fig 1D).

### Fitness proxies

Fitness proxies were calculated for 86 founders, 44 males and 42 females. We estimated the correlation with the reproductive values of 42 lineages that survived to 2018 and had no missing fitness data. In total, individuals included in the analysis produced 2,054 eggs of which 1,746 (85%) survived to hatching, 881 (43%) to fledging and 294 (14%) recruited into the breeding population.

The fitness proxies were all positively associated with the reproductive values (Fig. 2). De-lifed fitness, IGR and LRS for recruits were all statistically significantly correlated with the reproductive values. There were no statistically significant differences between the IGR, LRS and de-lifed fitness correlation coefficients. None of the other correlation estimates were statistically significant (Table 1). De-lifed fitness at ages 1 and 2 significantly correlated with reproductive value, but in older age classes the correlation estimate was not statistically significant (Figs 3A and 3B).

**Table 1:** Mean and standard deviation for short-term fitness proxies at different offspring stages, and de-lifed fitness at different ages.

|  | Mean | SD |
| --- | --- | --- |
| LRS Eggs | 23.78 | 18.02 |
| LRS Broods | 5.80 | 4.26 |
| LRS Hatchlings | 20.30 | 16.04 |
| LRS Fledglings | 10.24 | 7.85 |
| LRS Recruits | 3.42 | 3.36 |
| De-lifing | 0.01 | 0.03 |
| IGR eggs | 4.24 | 1.53 |
| IGR hatchlings | 3.71 | 1.37 |
| IGR fledglings | 2.43 | 0.99 |
| IGR recruits | 1.13 | 0.78 |
| De-lifing – Age 1 | 0.0003 | 0.0198 |
| De-lifing – Age 2 | 0.0056 | 0.0170 |
| De-lifing – Age 3 | 0.0027 | 0.0115 |
| De-lifing – Age 4 | 0.0023 | 0.0087 |
| De-lifing – Age 5 | 0.0033 | 0.0048 |
| De-lifing – Age 6 | 0.0005 | 0.0046 |

**Fig 2:**
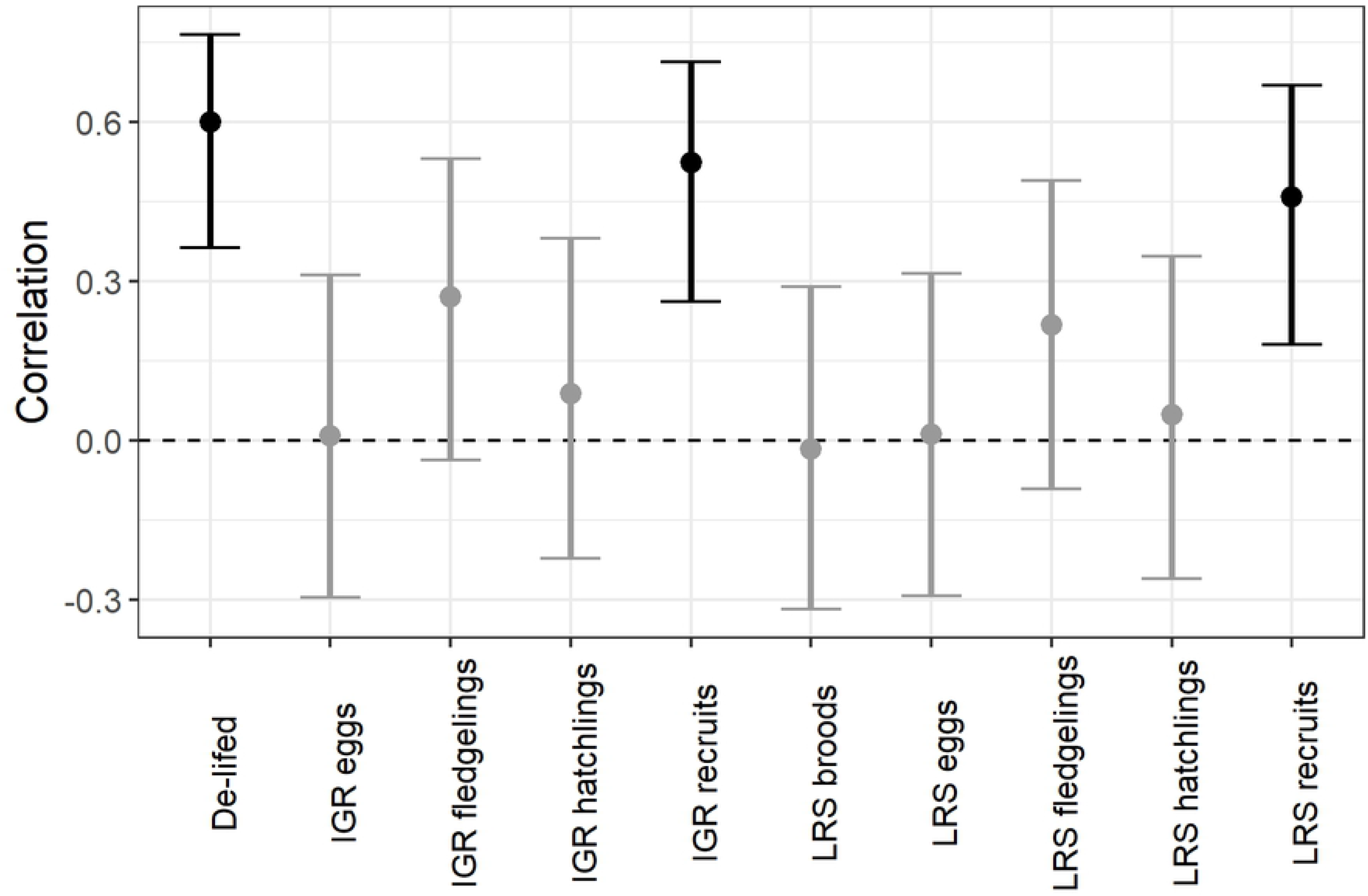
Correlation between each of the fitness proxies at different life stages with reproductive value. Error bars represent 95% confidence intervals. Black bars represent significant results, while light grey bars represent non-significant results.

**Fig 3:**
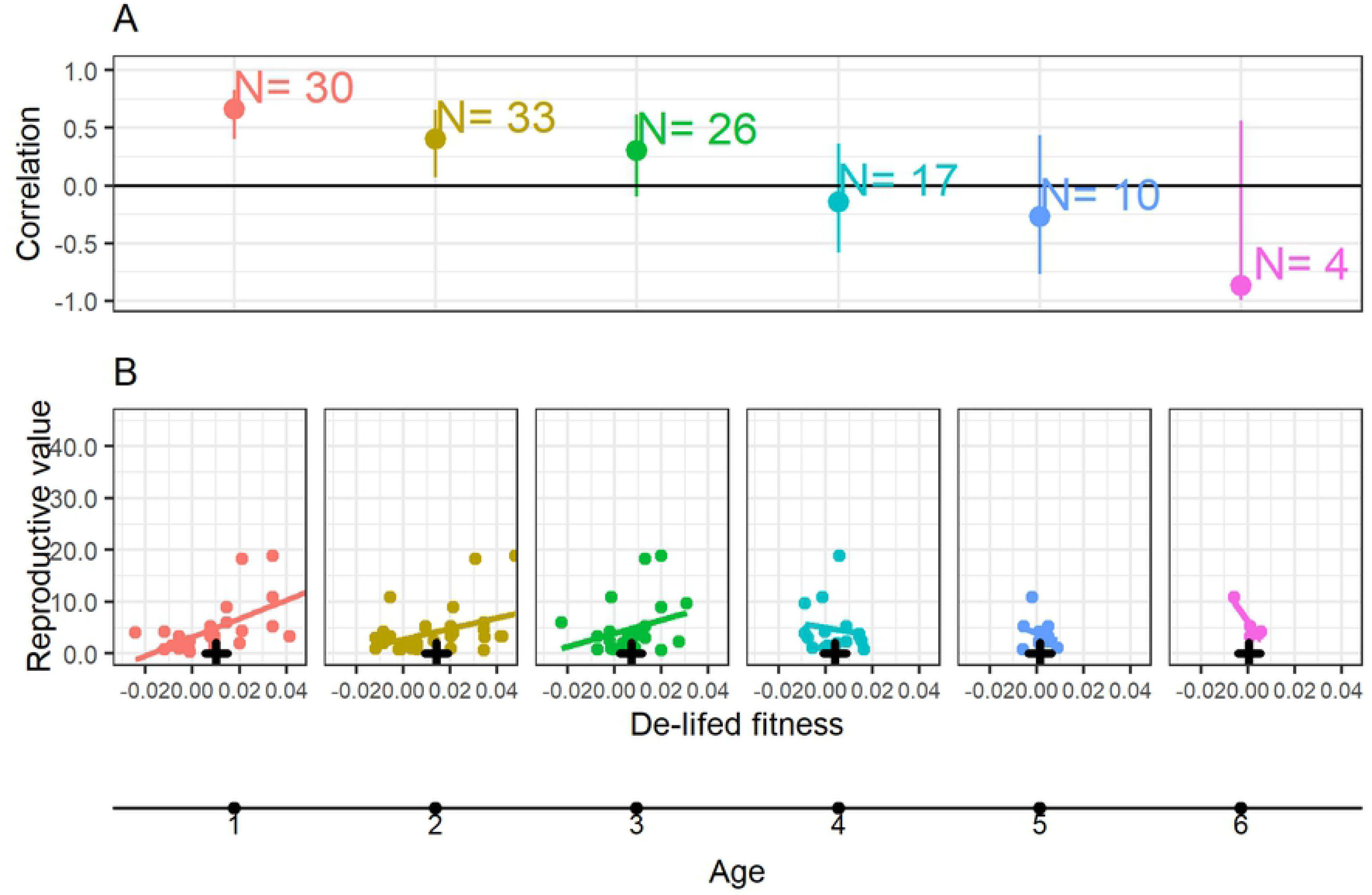
(A) Correlation between reproductive value and de-lifed fitness by age class (in years), with 95% confidence intervals. N represents the sample size. Older age classes have lower sample sizes because fewer individuals survive to that age. (B) Correlation between de-lifed fitness and reproductive value. The black cross represents mean de-lifed fitness for the respective age. Colours represent the same age class.

There was a significant positive relationship between lineage survival odds and de-lifed fitness, LRS at recruitment and fledgling stages, and IGR at recruitment (Table 2). The estimated slopes for the de-lifed fitness and LRS recruits were significantly higher than the slopes of IGR and LRS at fledgling stage as their 95% confidence intervals are non-overlapping, indicating that fitness measured at the recruitment stage for these two metrics is a better predictor of lineage survival. There were no statistically significant differences between IGR, LRS and de-lifed fitness at the same offspring stage (Table 2).

**Table 2:** Results of binary logistic regressions with lineage survival as a response variable. l-95% CI and u-95% CI are the lower and upper boundaries of the 95% confidence interval for the slope, respectively.

| Variable | Slope | l-95% CI | u-95% CI | p value |
| --- | --- | --- | --- | --- |
| LRS eggs | 0.42 | -0.07 | 0.95 | 0.101 |
| LRS hatchlings | 0.45 | -0.08 | 0.95 | 0.063 |
| LRS fledglings | 1.06 | 0.41 | 1.71 | 0.001* |
| LRS recruits | 3.26 | 1.73 | 4.74 | 0.001* |
| LRS broods | 0.47 | -0.02 | 0.96 | 0.056 |
| IGR eggs | 0.24 | -0.21 | 0.75 | 0.310 |
| IGR hatchlings | 0.35 | -0.09 | 0.87 | 0.167 |
| IGR fledglings | 1.06 | 0.49 | 1.71 | 0.002 |
| IGR recruits | 2.65 | 1.67 | 3.70 | 0.001* |
| De-lifed fitness | 3.01 | 1.74 | 4.32 | 0.001* |

## Discussion

We showed that fitness proxies measured at recruit stage correlates best with long-term reproductive values and lineage survival, while fitness proxies measured at earlier stages are less useful.

Similar to a study by (4) lineage survival is low. While there was no difference in the rate of lineage survival between native and introduced lineages, the introduced lineages had significantly higher reproductive values. This indicates that the introduced lineages might have a fitness advantage over the native ones. For lineages that survived to 2018, there was wide variation in survival probability and reproductive value. The survival probability of a lineage is associated with its reproductive value in that year (4,18). While several lineages died out every year prior to 2007, all lineages that survived the bottleneck in 2008 also survived the next 10 years to 2018. Lineage extinctions are expected to become less likely over the generations, as all founders with non-zero reproductive values become genealogical ancestors of all individuals in the future population after only a few generations (19,22). After a founder becomes an ancestor of all individuals in the current population, its lineage can only go extinct if the entire population goes extinct. During the 2008 bottleneck, the population size decreased significantly, shortening the time it took for all founders of persisting lineages to become the common ancestors of the current population members.

There was variation in reproductive value, with most lineages ranging from 0 to 10 but some reaching contributions of over four times that much. There was also large variation over time as lineages fluctuated. Lineage stabilisation is also visible in the pattern of lineage fluctuation through time after 2007, as the ranking of lineages based on reproductive value remains similar. The rapidly increasing correlation between reproductive value in the final year and each of the previous years also shows a pattern of stabilisation, as found in other studies too (4,18,34). Stabilisation is reached after 12 years, with the correlation exceeding 0.95 afterwards. Despite stabilisation, reproductive values fluctuated through time. As we examined reproductive values that are absolute rather than relative to population size, any change in population size is also reflected in the sum of the reproductive values the year before. The change in reproductive value occurs one year previously, because the estimates are based on reproducing offspring, which are only recognised in the next year and form the basis of next year’s population.

The fitness proxies based on the number of recruits outperformed all other fitness proxies in predicting reproductive values and lineage survival. Recruits are likely to be the best measure because they are adult individuals that reproduced, while other proxies include the uncertainty of survival to adulthood before reproduction even occurs. Given that sparrow offspring experience high rates of mortality, with only 14% of laid eggs successfully surviving to recruitment, mortality will have a big impact on reproductive values from short-term metrics measured at early offspring stages. For species with lower offspring mortality the age at which offspring are counted towards fitness may have less influence on the predictive power of short-term fitness metrics. While recruits are clearly the best predictor of long-term fitness, they are the most difficult to measure in most study systems, as it is rarely possible to monitor all offspring until their first reproduction. This highlights the importance of long-term isolated island population studies (35), as only in such studies is it possible to accurately estimate the number of genetic recruits that an individual produced.

We found no differences in the performance of de-lifed fitness, IGR or LRS in predicting reproductive values or lineage survival. A previous study on Ural owls (*Strix uralensis*) and collared flycatchers (*Ficedula albicollis*) found that LRS performed significantly better than IGR at fledgling stage in predicting reproductive values, while they both performed similarly at recruitment (10). The correlation between reproductive value and different fitness proxies at recruit stage was of similar strength as discovered in previous studies (4,10).

In this study, annual de-lifed fitness at ages 1 and 2 were correlated with the reproductive values, but not at later ages. The correlation at age 1 with reproductive value was similar to that for lifetime de-lifed fitness, indicating that reproductive success in the first adult year may be sufficient to provide a good prediction of long-term fitness. Hence, individual reproductive performance in the first year may be an important proxy for an individual’s fitness.

There is, however, considerable variation that is not explained by the fitness metrics. A strong correlation between a short-term fitness metric and the reproductive value measured two decades later, during which the population has been exposed to varying environmental conditions and population fluctuations, is unlikely. The strength of the correlation will also depend on the additive genetic variance and heritability of reproductive success (4). In particular, in our population annual fitness is somewhat heritable (36), and there has been significant demographic stochasticity in our population for which LRS and IGR metrics tested here were not designed (37).

The underlying theoretical results about reproductive value were derived under the assumption of diploid Wright-Fisher population and weak selection (18). The Lundy sparrow population might not meet these assumptions, as there could be undetected strong selection and non-random mating. The sparrow population can therefore be used to test theoretical predictions on real data but could lead to erroneous conclusions if assumptions are severely breached. Despite that, reproductive values can lead to new insights about natural selection and evolutionary outcomes, such as inbreeding, lineage introgression or cohort effects (2,4,34). Particularly in the presence of environmental, social, or demographic interactions, such as those occurring in any wild population, studying fitness across an entire lineage by examining reproductive values can potentially lead to better estimation of evolutionary outcomes for a certain allele (38).

In conclusion, by using reproductive values as a measure of long-term individual fitness we have shown that recruits, rather than earlier offspring stages, best predict reproductive values. Additionally, annual fitness measured in the first reproductive season is an equally good predictor of fitness as lifetime fitness measures. We therefore suggest that future studies should measure short-term fitness at higher offspring ages to better capture long-term fitness.

## Author Contributions

Conceptualization: Vita Živa Alif, Julia Schroeder, Terry Burke

Data Curation: Jamie Dunning, Janet Chik, Julia Schroeder, Terry Burke

Formal Analysis: Vita Živa Alif

Funding acquisition: Terry Burke, Julia Schroeder

Investigation: Vita Živa Alif

Project administration: Terry Burke, Julia Schroeder

Resources: Terry Burke, Julia Schroeder

Software: Vita Živa Alif

Supervision: Julia Schroeder

Validation: Vita Živa Alif

Visualisation: Vita Živa Alif

Writing – original draft: Vita Živa Alif

Writing – review & editing: Vita Živa Alif, Julia Schroeder, Jamie Dunning, Janet Chik, Terry Burke

## References

1. Fisher RA. The genetical theory of natural selection. [Internet]. Oxford: Clarendon Press; 1930. 302 p. Available from: https://www.biodiversitylibrary.org/bibliography/27468

2. Grafen A. The Price equation and reproductive value. Philos Trans R Soc B Biol Sci. 2020 Apr 27;375(1797):20190356.

3. Orr HA. Fitness and its role in evolutionary genetics. Nat Rev Genet. 2009 Aug;10(8):531–9.

4. Reid JM, Nietlisbach P, Wolak ME, Keller LF, Arcese P. Individuals’ expected genetic contributions to future generations, reproductive value, and short-term metrics of fitness in free-living song sparrows (Melospiza melodia). Evol Lett. 2019;3(3):271–85.

5. Saether B-E, Engen S. The concept of fitness in fluctuating environments. Trends Ecol Evol. 2015;30(5):9.

6. Brommer JE, Merilä J, Kokko H. Reproductive timing and individual fitness. Ecol Lett. 2002;5(6):802–10.

7. Dobson FS, Murie J, Viblanc V. Fitness Estimation for Ecological Studies: An Evaluation in Columbian Ground Squirrels. Front Ecol Evol [Internet]. 2020 [cited 2020 Nov 11];8. Available from: https://hal.archives-ouvertes.fr/hal-02903692

8. Coulson T, Benton T g, Lundberg P, Dall S r. x, Kendall B e, Gaillard J-M. Estimating individual contributions to population growth: evolutionary fitness in ecological time. Proc R Soc B Biol Sci. 2006 Mar 7;273(1586):547–55.

9. Dobson FS, Lane JE, Low M, Murie JO. Fitness implications of seasonal climate variation in Columbian ground squirrels. Ecol Evol. 2016;6(16):5614–22.

10. Brommer JE, Gustafsson L, Pietiäinen H, Merilä J. Single-generation estimates of individual fitness as proxies for long-term genetic contribution. Am Nat. 2004 Apr 1;163(4):505–17.

11. Moorad JA. Individual fitness and phenotypic selection in age-structured populations with constant growth rates. Ecology. 2014;95(4):1087–95.

12. Roff DA. Defining fitness in evolutionary models. J Genet. 2008 Dec 1;87(4):339–48.

13. McGraw JB, Caswell H. Estimation of Individual Fitness from Life-History Data. Am Nat [Internet]. 1996 [cited 2020 Nov 17]; Available from: https://www.journals.uchicago.edu/doi/abs/10.1086/285839

14. Rubach K, Dobson FS, Zinner B, Murie J, Viblanc V. Comparing fitness measures and the influence of age of first reproduction in Columbian ground squirrels. J Mammal [Internet]. 2020 [cited 2020 Dec 8]; Available from: https://hal.archives-ouvertes.fr/hal-02916845

15. Grafen A. On the uses of data on lifetime reproductive success. In: Clutton-Brock TH, editor. Reproductive success. Chicago, Illinois, USA: University of Chicago Press; 1988. p. 454–71.

16. Käär P, Jokela J. Natural selection on age–specific fertilities in human females: comparison of individual–level fitness measures. Proc R Soc Lond B Biol Sci. 1998 Dec 22;265(1413):2415–20.

17. Wolf JB, Wade MJ. On the assignment of fitness to parents and offspring: whose fitness is it and when does it matter? J Evol Biol. 2001;14(2):347–56.

18. Barton NH, Etheridge AM. The relation between reproductive value and genetic contribution. Genetics. 2011 Aug;188(4):953–73.

19. Chang JT. Recent common ancestors of all present-day individuals. Adv Appl Probab. 1999;31(4):1002–26.

20. Chen N, Juric I, Cosgrove EJ, Bowman R, Fitzpatrick JW, Schoech SJ, et al. Allele frequency dynamics in a pedigreed natural population. Proc Natl Acad Sci. 2019 Feb 5;116(6):2158–64.

21. Hill WG. Variation in genetic identity within kinships. Heredity. 1993 Dec;71(6):652–3.

22. Gravel S, Steel M. The existence and abundance of ghost ancestors in biparental populations. Theor Popul Biol. 2015 May 1;101:47–53.

23. Grafen A. A theory of Fisher’s reproductive value. J Math Biol. 2006 Jul;53(1):15–60.

24. Schroeder J, Nakagawa S, Rees M, Mannarelli M-E, Burke T. Reduced fitness in progeny from old parents in a natural population. Proc Natl Acad Sci. 2015 Mar 31;112(13):4021–5.

25. Kozłowski J. Measuring fitness in life-history studies. Trends Ecol Evol. 1993 Mar;8(3):84–5.

26. Ockendon N, Griffith SC, Burke T. Extrapair paternity in an insular population of house sparrows after the experimental introduction of individuals from the mainland. Behav Ecol. 2009 Mar 1;20(2):305–12.

27. Schroeder J, Cleasby IR, Nakagawa S, Ockendon N, Burke T. No evidence for adverse effects on fitness of fitting passive integrated transponders (PITs) in wild house sparrows Passer domesticus. J Avian Biol. 2011;42(3):271–5.

28. Simons MJP, Winney I, Nakagawa S, Burke T, Schroeder J. Limited catching bias in a wild population of birds with near-complete census information. Ecol Evol. 2015;5(16):3500–6.

29. MacCluer JW, VandeBerg JL, Read B, Ryder OA. Pedigree analysis by computer simulation. Zoo Biol. 1986;5:147–60.

30. Wolak ME. nadiv : an R package to create relatedness matrices for estimating non-additive genetic variances in animal models. Methods Ecol Evol. 2012;3(5):792–6.

31. Gratten J, Wilson AJ, McRae AF, Beraldi D, Visscher PM, Pemberton JM, et al. A Localized Negative Genetic Correlation Constrains Microevolution of Coat Color in Wild Sheep. Science. 2008 Jan 18;319(5861):318–20.

32. R Core Team. R: A Language and Environment for Statistical Computing. Vienna, Austria: R Foundation for Statistical Computing; 2020.

33. Hadfield JD. MCMC Methods for Multi-Response Generalized Linear Mixed Models: The MCMCglmm R Package. J Stat Softw [Internet]. 2010 [cited 2021 Jan 23];33(2). Available from: http://www.jstatsoft.org/v33/i02/

34. Hunter DC, Pemberton JM, Pilkington JG, Morrissey MB. Pedigree-based estimation of reproductive value. J Hered. 2019 Jul 1;110(4):433–44.

35. Clutton-Brock T, Sheldon BC. Individuals and populations: the role of long-term, individual-based studies of animals in ecology and evolutionary biology. Trends Ecol Evol. 2010 Oct 1;25(10):562–73.

36. Schroeder J, Burke T, Mannarelli M-E, Dawson DA, Nakagawa S. Maternal effects and heritability of annual productivity: Maternal effects of annual productivity. J Evol Biol. 2012 Jan;25(1):149–56.

37. Engen S, Lande R, Sæther B, Dobson FS. Reproductive value and the stochastic demography of age-structured populations. Am Nat. 2009 Dec;174(6):795–804.

38. Graves CJ, Weinreich DM. Variability in Fitness Effects Can Preclude Selection of the Fittest. Annu Rev Ecol Evol Syst. 2017;48(1):399–417.

